## Supporting Information for "Phase separation driven by interchangeable properties in the intrinsically disordered regions of protein paralogs"

S.-H. Chiu et al.

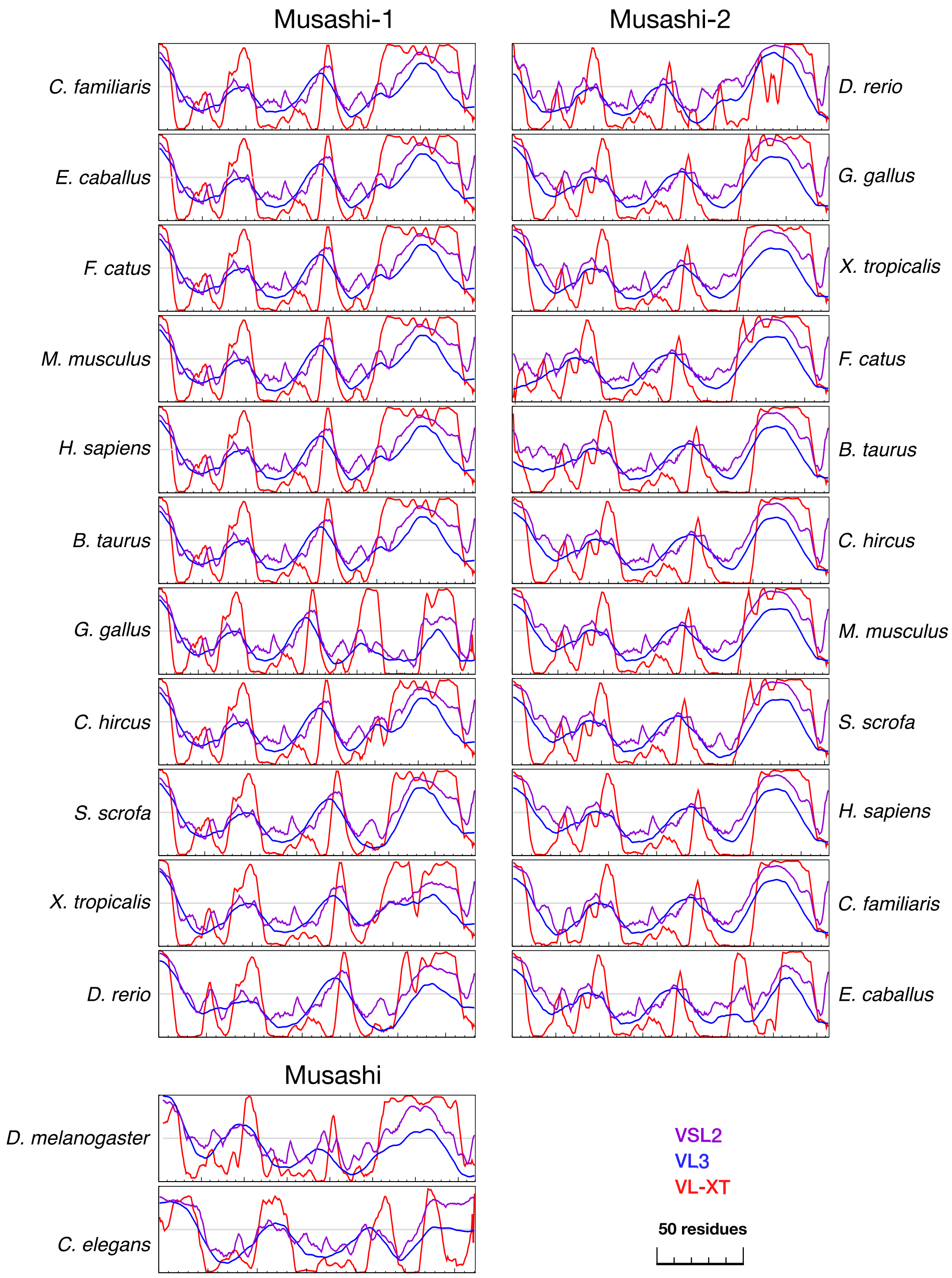


**Figure S1.** Structural disorder predictions for Musashi proteins. Three algorithms: VSL2 (purple), VL3 (blue), and VL-XT (red) were used. The species are in the same order as in fig. 1*A*.

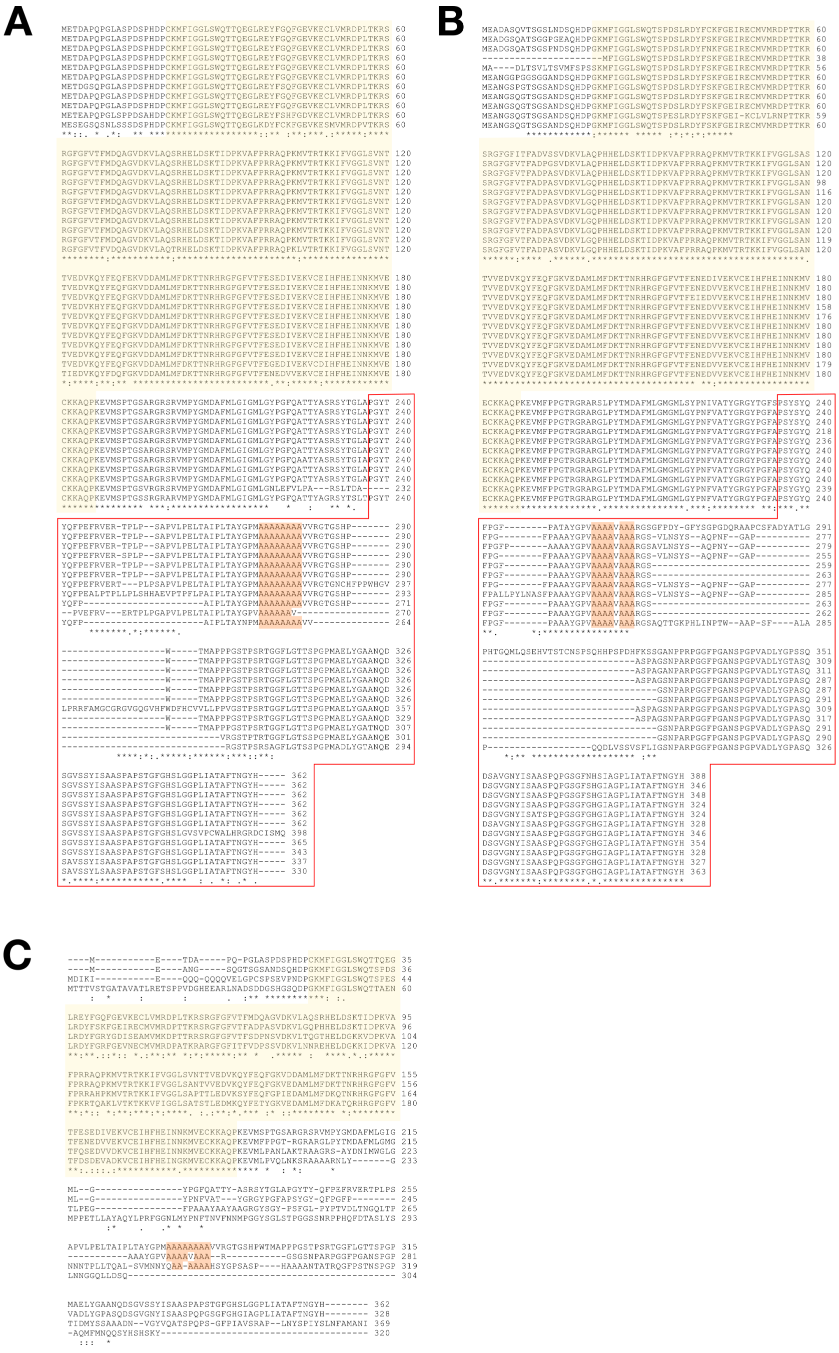


**Figure S2.** Sequence alignment of Musashi proteins. (*A*) Musashi-1 (*B*) Musashi-2 in vertebrates (with species in the same order as in fig. 1*B*), and (*C*) human Musashi-1 and -2 with nematode and fruit fly orthologs. The RRMs are highlighted in yellow and the red boxes indicate the IDRs (according to the definition in the main text and alignment to human orthologs). The polyalanine region is highlighted in orange.

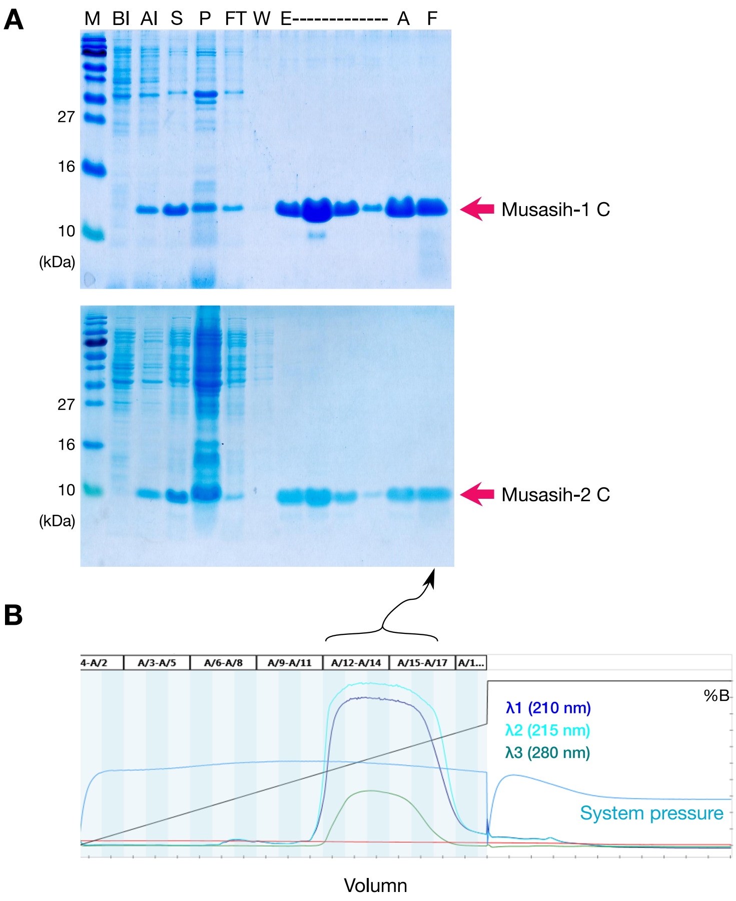


**Figure S3.** Examples of protein purification results. (*A*) SDS-PAGE gels of Musashi-1 (top) and Musashi-2 (bottom). M: marker; BI/AI: before/after IPTG induction; S/P: supernatant/pellet of lysed cell; FT/W/E: flow-through/wash-through/elution of the IMAC purification; A: acidified sample before loading into the C4 column; F: the final sample obtained from the HPLC column, purified and then lyophilized. (*B*) A typical HPLC elution profile. The fractions collected (indicated with curly bracket) were lyophilized before use.

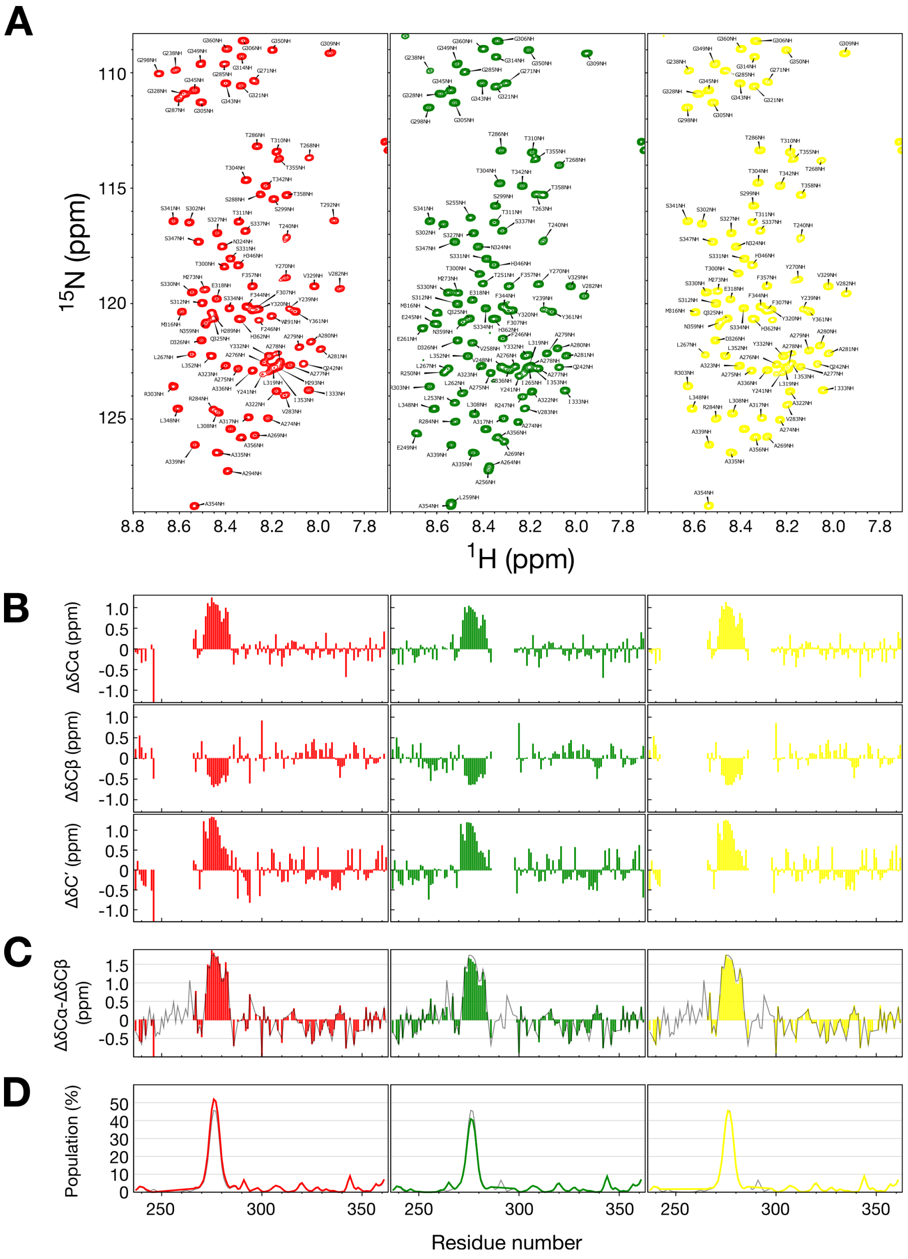


**Figure S4.** NMR analysis of the Msi-1C deletion variants. (*A*) HSQC spectra and chemical shift assignments, ΔSeq1 (red), ΔSeq2 (green), and ΔSeq1ΔSeq2 (yellow). (*B*) Cα, Cβ, and C' secondary chemical shifts, (*C*) secondary chemical shift differences between Cα and Cβ atoms (to eliminate chemical shift referencing errors), with the results for wild-type Msi-1C shown in gray for comparison; (*D*) α-helix populations calculated with the δ2D algorithm (from H, N, Cα, Cβ, and C' chemical shifts) with the results for the wild type shown in gray. The differences between the variants and the wild type for panels (*C*) and (*D*) are shown in fig. 3*F and G* in the main text.

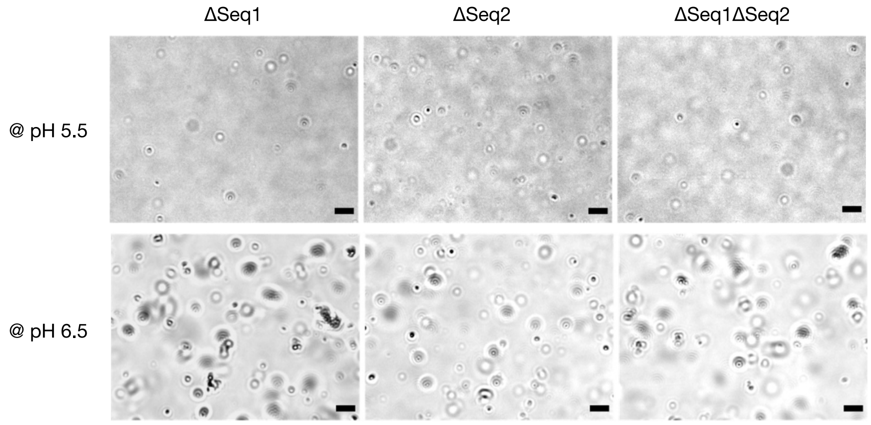


**Figure S5.** Optical micrographs of the condensates observed in the Msi-1C deletion constructs. ΔSeq1, ΔSeq2, and ΔSeq1ΔSeq2 at pH 5.5 and 6.5 are shown. Scale bar: 10 μm.

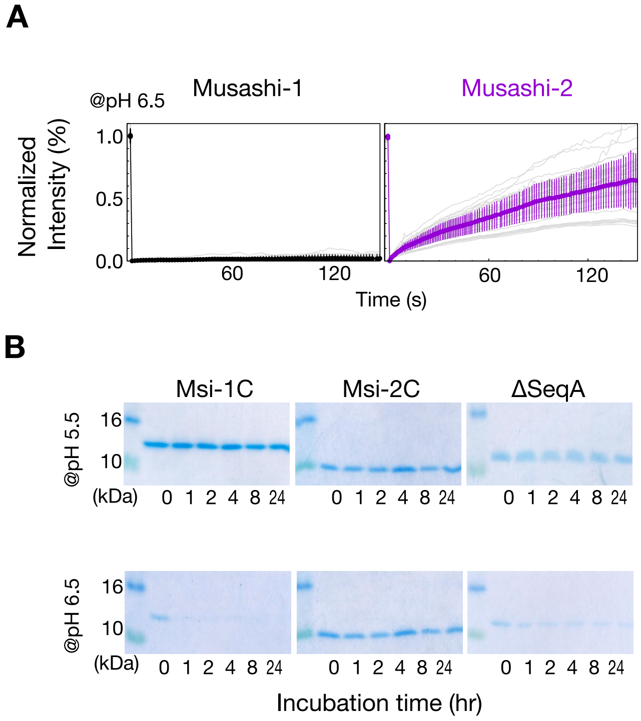


**Figure S6.** FRAP and SDS-PAGE analysis. (*A*) The FRAP analysis of Musashi-1 and Musashi-2 at pH 6.5. The colored lines represent the mean ± standard deviation and the gray lines are individual recovery profiles. (*B*) SDS-PAGE gels of precipitation assays of the supernatants of centrifuged Msi-1C, Msi-2C, and ΔSeqA samples after different incubation times at pH 5.5 or pH 6.5.

**A**

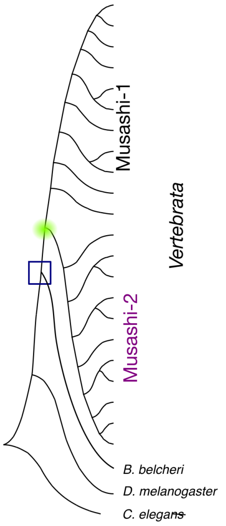


**B**

```
fruit fly  MDIKIEQQQQQQVELGPCSPSEVPNDPGKMFIGGLSWQTSPESLRDYFGRYGDISEAMV 60
lancelet   --------MDGQSQQTSANSPSEEANDPGKMFIGGLSWQTTADSLREYFCKYGEIKECMV 52
                   : *. : .  ****  **************: :***:** :**:*.*.**
           MKDPTTRRSRGFGFVTFSDPNSVDKVLTQGTHELDGKKVDPKVAFPRRAHPKMVTRTKKI 120
           MKDPVTRRSRGFGFVTFRDPASVDKVLAVSKHELDSKTIDPKVAFPRRAQPKMVTRTKKI 112
           ****.************ ** ******: ..****.*.:**********:**********
           FVGGLSAPTTLEDVKSYFEQFGPIEDAMLMFDKQTNRHRGFGFVTFQSEDVVDKVCEIHF 180
           FVGGLSANTVLEDVKNYFGHYGKIEDAMLMFDKQTNRHRGFGFVTFESEDIVDKVCDIHF 172
           ******* *.*****.** ::* **********************:***:*****:***
           HEINNKMVECKKAQPKEVMLPANLAKTRAAGRSAY--------DNIMWGLGTLPEGFPAA 232
           HEINNKMVECKKAQPKEVMFPPGSARGRARGAFGTWPDLGESFLYLLHPEIVREDGIQAG 232
           ********************:* . *: ** *  .            ::    .  :*: *.
           AYAAYAAGRGYSGYPSFGLPYPTVDLTN---G----QL-TPNNNTPLLTQALSVMNNYQA 284
           VYPNFAYGRGYPFAPGYGYGFPESGHFVFPTGFPAAAYGPPGRASPRGARGRAGIMGYPG 292
           .*  :* ****   *.:*  :*   .       *         *.. :*  ::. : : .* .
           AAAAAHSYGPSASPHAAAANTATRQGFPSTNSPGPTIDMYSSAAAD----NVGYVQATSP 340
           TA-VLNSYAT--QGFAPPASPANSRGFPPANSPGPI-DMYGGTTSQADTVSSYVAQAASP 348
           :* . :**.    . .*  *. *. :*** :*****  ***..::::    .   .**:**
           QPSGFPIAVSRAPL--NYSPIYS--LNFAMANI 369
           QPSGFSIAPSSGFGNSNYAGMVGNPRNF----- 376
           ***** ** * .    **: : .   **
```

**C**

**Musashi-1**

```
human        METDAPQ-PGLASPDSPHDPCKMFIGGLSWQTTQEGLREYFGQFGEVKECLVMRDPLTKR 59
ghost shark  MEADANQANSVLSADSQHDPCKMFVGGLSWQTTQEGLKEYFCKFGDVKECMVMRDPVTKR 60
             **:** *  .: * ** *******:***********:***  :**:****:*****:***
             SRGFGFVTFMDQAGVDKVLAQSRHELDSKTIDPKVAFPRRAQPKMVTRTKKIFVGGLSVN 119
             SRGFGFVTFVDQTGVDKVLAQPRHELDSKTIDPKVAFPRRAQPKMVTRTKKIFVGGLSVN 120
             *********:**:******** **************************************
             TTVEDVKQYFEQFGKVDDAMLMFDKTTNRHRGFGFVTFESEDIVEKVCEIHFHEINNKMV 179
             TTVEDVKQYFDQFGKVDDAMLMFDKTTNRHRGFGFVTFESEDIVEKVCEIHFHEINNKMV 180
             **********:*************************************************
             ECKKAQPKEVMSPTGSARGRSRVMPYGMDAFMLGIGMLGYPGFQATTYASRSYTGLAPGY 239
             ECKKAQPKEVMSPTGTARGRSRVMPYGMDAFMLGIGMLGYPGFQAASYASRGYTGLAPGY 240
             ***************:****************************::****.********
             TYQFPEFRVERT--PLP-----------------SAPVLP--ELTAIPLTAYGPMAAAAA 278
             TYQFPGPFLLPPTATLPSSAGRAAQSALQTSQWPPSVLLQDWPTGAIPLTAYGPMAAAAA 300
             *****    :      **                 : :*      **************
             AAAVVRGTGSHPWTMAPPPGSTPSRTGGFLGTTSPGPMAELYGAANQDSGVSSYISAASP 338
             AA-VVRGT-----------GSTPTRTGGFLGTTSPAPMADLYGTANQDSGVSSYISAASP 348
             * ******           ****:**********.***:***:****************
             APSTGFGHSLGGPLIATAFTNGYH 362
             APSTGFGHSLGGPLIATAFTNGYH 372
             ************************
```

**Musashi-2**

```
MEANGSQGTSGSANDSQHDPGKMFIGGLSWQTSPDSLRDYFSKFGEIRECMVMRDPTTKR 60
MEADGSQATSGSPSDSQHDPGKMFIGGLSWQTSPDSLRDYFSKFGEIRECMVMRDPTTKR 60
***:***.****  .*********************************************
SRGFGFVTFADPASVDKVLGQPHHELDSKTIDPKVAFPRRAQPKMVTRTKKIFVGGLSAN 120
SRGFGFVTFADPASVDKVLAQPHHELDSKTIDPKVAFPRRAQPKMVTRTKKIFVGGLSAN 120
*******************.****************************************
TVVEDVKQYFEQFGKVEDAMLMFDKTTNRHRGFGFVTFENEDVVEKVCEIHFHEINNKMV 180
TVVDDVKQYFEQFGKIEDAMLMFDKTTNRHRGFGFVTFENEEVVEKVCEIHFHEINNKMV 180
***:***********:************************:*******************
ECKKAQPKEVMFPPGTRGRARGLPYTMDAFMLGMGMLGYPNFVATYGRGYPGFAPSYGYQ 240
ECKKAQPKEVMFPPGTRGRARGLPYTMDAFMLGMGMLGYPNFVATYGRGYPGFAPSYSYQ 240
*********************************************************.**
FPGFPAAAYGPVAAAAVAA----------------------------------------- 259
FPAFPAAAYGPVAAAAAAARGSGFPDYGFYSSPGDQRASVSFADYGSMGLRAAQMLRSEH 300
**.*************.**
-----------------------------------ARGSGSNPARPGGFPGANSPGPVAD 284
APQASSPSLHHLSSPDQFKNPVLNSYTAQPNFGGPASPAGTNPARPGGFPGANSPGPVAD 360
                                   *   :*:******************
LYGPASQDSGVGNYISAASPQPGSGFGHGIAGPLIATAFTNGYH 328
IYGTASQDSGVGNYMSATSPQPGSGFGHSIGGPLIATAFTNGYH 404
:** **********:**:**********.*.*************
```

**Figure S7.** Sequence analysis of primitive chordates. (*A*) Phylogenetic tree of the Musashi family as shown in fig. 1*A* with the additional lineages analyzed in this figure indicated as the blue box and green circle. (*B*) Sequence alignment of fruit fly (*D. melanogaster*) and lancelet (*B. belcheri*; UniProt entry: A0A6P4YVJ9) Musashi protein. The RNA recognition motifs (as predicted by PROSITE) are indicated in yellow. (*C*) Sequence alignment of human and ghost shark (*C. milii*) Musashi-1 (left, UniProt entry: V9KSD1) and Musashi-2 (right, UniProt entry: V9KVG4). The polyalanine tracts are highlighted in orange.

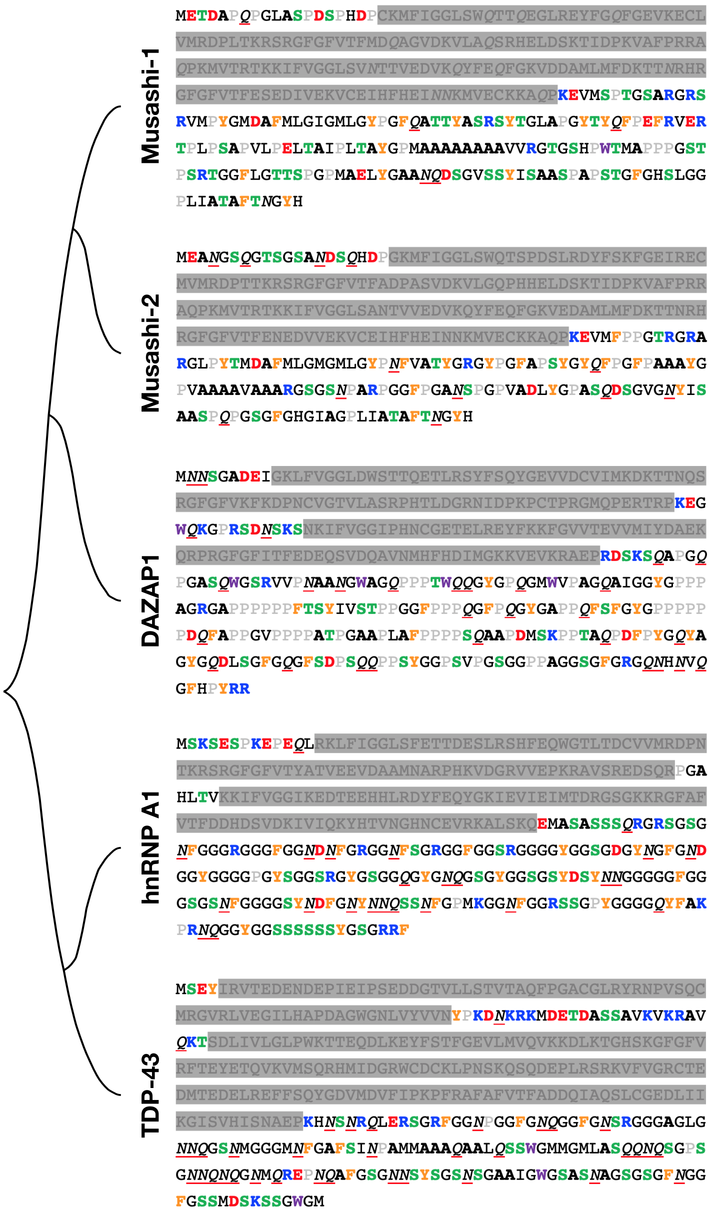


**Figure 8.** Primary sequences of Musashi paralogs. The structured domains (as defined by PROSITE) are shaded in gray. The amino acids are color coded based on their physical properties (positive charge, blue; negative charge, red; F/Y, yellow; W, purple; S/T (potential phosphorylation site for the addition of negative charges), green; P, grey; A, bold black; Q/N, red underlined italic)

**Table S1.** Genes used in this study.

| Entry | Protein (submitted name) | Gene | Organism |
|---|---|---|---|
| O43347 | Rbp Musashi homolog 1 | MSI1 | *Homo sapiens* (human) |
| Q96DH6 | Rbp Musashi homolog 2 | MSI2 | *Homo sapiens* (human) |
| Q61474 | Rbp Musashi homolog 1 | MSI1 | *Mus musculus* (mouse) |
| Q920Q6 | Rbp Musashi homolog 2 | MSI2 | *Mus musculus* (mouse) |
| E2RK48 | Musashi Rbp 1 | MSI1 | *Canis lupus familiaris* (dog) |
| A0A5F4BU43 | Musashi Rbp 2 | MSI2 | *Canis lupus familiaris* (dog) |
| A0A337SLM7 | Musashi Rbp 1 | MSI1 | *Felis catus* (cat) |
| A0A337SE56 | Uncharacterized protein | MSI2 | *Felis catus* (cat) |
| F6SHE7 | Musashi Rbp 1 | MSI1 | *Equus caballus* (horse) |
| F7AKB0 | Musashi Rbp 2 | MSI2 | *Equus caballus* (horse) |
| A2PYH9 | Musashi Rbp 1 | MSI1 | *Bos taurus* (cattle) |
| A0A3Q1M610 | Uncharacterized protein | MSI2 | *Bos taurus* (cattle) |
| A0A452FXY9 | Uncharacterized protein | MSI1 | *Capra hircus* (goat) |
| A0A452EW36 | Uncharacterized protein | MSI2 | *Capra hircus* (goat) |
| I3LPG0 | Musashi Rbp 1 | MSI1 | *Sus scrofa* (pig) |
| A0A287AG72 | Musashi Rbp 2 | MSI2 | *Sus scrofa* (pig) |
| A0A3Q2UHP1 | Uncharacterized protein | MSI1 | *Gallus gallus* (chicken) |
| E1C1R8 | Uncharacterized protein | MSI2 | *Gallus gallus* (chicken) |
| A0A6I8QUP6 | Musashi Rbp 1 | MSI1 | *Xenopus tropicalis* (frog) |
| A0A6I8SE33 | Uncharacterized protein | MSI2 | X *Xenopus tropicalis* (frog) |
| Q5BKV4 | Msi1 protein | MSI1b | *Danio rerio* (zebrafish) |
| Q7ZW10 | Msi2 protein | MSI2b | *Danio rerio* (zebrafish) |
| Q9VVE5 | Rbp Musashi homolog Rbp6 | Rbp6 | *Drosophila melanogaster* (fruit fly) |
| G5EFS2 | MuSashI (Fly neural) family | Msi-1 | *Caenorhabditis elegans* (nematode) |

**Table S2.** Frequency and percentage of each amino acid in the RRMs and IDRs of Musashi-1 and Musashi-2.

| | Msi-1 RRMs | | Msi-2 RRMs | | Msi-1 IDRs | | Msi-2 IDRs | |
|---|---|---|---|---|---|---|---|---|
| aa | count | % | count | % | count | % | count | % |
| A | 138 | **5.3** | 141 | **5.5** | 215 | **15.9** | 242 | **19.7** |
| R | 153 | **5.9** | 153 | **6** | 43 | **3.2** | 23 | **1.9** |
| N | 47 | **1.8** | 80 | **3.1** | 23 | **1.7** | 58 | **4.7** |
| D | 157 | **6.1** | 151 | **5.9** | 12 | **0.9** | 27 | **2.2** |
| C | 44 | **1.7** | 34 | **1.3** | 5 | **0.4** | 2 | **0.2** |
| Q | 117 | **4.5** | 82 | **3.2** | 23 | **1.7** | 45 | **3.7** |
| E | 165 | **6.4** | 153 | **6** | 38 | **2.8** | 1 | **0.1** |
| G | 238 | **9.2** | 253 | **9.9** | 172 | **12.7** | 200 | **16.3** |
| H | 57 | **2.2** | 64 | **2.5** | 36 | **2.7** | 28 | **2.3** |
| I | 78 | **3.0** | 70 | **2.7** | 32 | **2.4** | 35 | **2.8** |
| L | 132 | **5.1** | 104 | **4.1** | 88 | **6.5** | 36 | **2.9** |
| K | 189 | **7.3** | 187 | **7.3** | 0 | **0** | 2 | **0.2** |
| M | 142 | **5.5** | 141 | **5.5** | 31 | **2.3** | 1 | **0.1** |
| F | 187 | **7.2** | 209 | **8.2** | 55 | **4.1** | 65 | **5.3** |
| P | 139 | **5.4** | 155 | **6.1** | 184 | **13.6** | 169 | **13.8** |
| S | 148 | **5.7** | 128 | **5** | 132 | **9.8** | 133 | **10.8** |
| T | 185 | **7.1** | 165 | **6.5** | 130 | **9.6** | 33 | **2.7** |
| W | 11 | **0.4** | 11 | **0.4** | 11 | **0.8** | 1 | **0.1** |
| Y | 63 | **2.4** | 66 | **2.6** | 63 | **4.7** | 76 | **6.2** |
| V | 202 | **7.8** | 200 | **7.9** | 60 | **4.4** | 52 | **4.2** |

**Table S3**. Primers used in this study.

| Constructs | Primers (5'–3') |
|---|---|
| MSI-1[237-362] | *Fw*-GGAGATATACATATGCCTGGCTACACCTAC |
| | *Rv*-GTAGGTGTAGCCAGGCATATGTATATCTCCTTCTTAAAGTT |
| MSI-1-ΔSeq1 | Fw-GAATTCCCTCTCACTGCCTAC |
| | *Rv*-GAGAGGGAATTCGGGGAACTGG |
| MSI-1-ΔSeq2 | *Fw*-GGGACAGGTTCGACTCCCAG |
| | *Rv*-CGAACCTGTCCCTCGAACCAC |
| MSI-1-ΔSeq1Seq2 | The same as ΔSeq1 and ΔSeq2 |
| MSI-1-ΔSeqA | *Fw*-GCCATTGGCTCTCACCCCTGG |
| | *Rv*-AGAGCCAATGGCTGTAAGCTC |
| MSI-2[235-328] | *Fw*-CCAAGCTATGGCTATCAG |
| | *Rv*-ATAGCCATAGCTTGGCATATGTATATCTCCTTCTTAAAGTT |
